# Stress activity is not predictive of coping style in North American red squirrels

**DOI:** 10.1101/465187

**Authors:** Sarah E. Westrick, Freya van Kesteren, Rupert Palme, Rudy Boonstra, Jeffery E. Lane, Stan Boutin, Andrew G. McAdam, Ben Dantzer

## Abstract

Individuals vary in their behavioral and physiological responses to environmental changes. These behavioral responses are often described as ‘coping styles’ along a proactive-reactive continuum. Studies in laboratory populations often, but not always, find that behavioral responses and physiological responses to stressors covary, where more proactive (more aggressive and active) individuals have a lower physiological stress response, specifically as measured by hypothalamic-pituitary-adrenal (HPA) axis activity. These studies support the possibility of hormonal pleiotropy underlying the presentation of behaviors that make up the proactive-reactive phenotype. However, recent research in wild populations is equivocal, with some studies reporting the same pattern as found in many controlled laboratory studies, whereas others do not. We tested the hypothesis that physiological and behavioral stress responses are correlated in wild adult North American red squirrels (*Tamiasciurus hudsonicus*). We used fecal cortisol metabolites (FCMs) as a non-invasive, integrated estimate of circulating glucocorticoids for our measurement of HPA axis activity. We found that FCM concentrations were not correlated with three measures of behavioral coping styles (activity, aggression, and docility) among individuals. This does not support the hypothesis that hormonal pleiotropy underlies a proactive-reactive continuum of coping styles. Instead, our results support the “two-tier” hypothesis that behavioral and physiological stress responses are independent and uncorrelated traits among individuals in wild populations that experience naturally varying environments rather than controlled environments. If also found in other studies, this may alter our predictions about the evolutionary consequences of behavioral and endocrine coping styles in free-living animals.

**Significance Statement:** Individuals vary in how they respond to stressors through behavior and physiology, but we find the two responses are independent in wild animals. Many laboratory studies find links between the behavioral and physiological stress responses, however studies conducted with wild populations are less conclusive. In wild North American red squirrels, independence between the physiological response and behavioral response may allow adaptive responses to a changing environment without pleiotropic constraint.

## Introduction

Organisms can respond to fluctuating environmental challenges and aversive stimuli both through behavioral responses and physiological stress responses. Laboratory studies often find these responses to be associated with one another (but see Steimer and Driscoll 2003; Koolhaas et al. 2007). In behavioral ecology and behavioral neuroscience, ‘coping styles’ have been recognized as one method of categorizing behavioral reactions to environmental challenges and stressors. Coping styles refer to a consistent set of behavioral responses to a stressor (Gosling 2001; Réale et al. 2007; Koolhaas et al. 2010; Stamps and Groothuis 2010). Furthermore, the suite of behaviors that make up an individual’s coping style is theorized to be mediated by hormones that exert pleiotropic actions (Koolhaas et al. 1999; McGlothlin and Ketterson 2008).

This unidimensional model has been repeatedly supported by studies describing how the hypothalamic-pituitary-adrenal (HPA) axis mediates coping styles (Koolhaas et al. 1999). Many of these studies have used selected lines, or have been done under controlled conditions in the laboratory. The conclusion from this model is that the behavioral stress response and physiological stress response run along the same axis. This hypothesis suggests a unidimensional response along a proactive-reactive continuum, where ‘proactive’ individuals are highly aggressive, highly active, and exhibit lower HPA axis reactivity and activity compared to ‘reactive’ individuals (Koolhaas et al. 1999; Cockrem 2007; Carere et al. 2010). The vast majority of these studies have been conducted using laboratory animals or wild animals selected for specific behavioral phenotypes, producing individuals at the extremes of this behavioral continuum. For example, in wild Great Tits (*Parus major*) lines selected for divergent personality types show the predicted unidimensional relationship between behavioral and stress responses in that more proactive birds exhibited lower HPA axis reactivity in response to capture and restraint (Baugh et al. 2012), and lower baseline corticosterone metabolites (Stöwe et al. 2010).

As more empirical studies are testing these models, the results from studies in the wild have been equivocal. Whereas there is some support for the unidimensional model in wild animals (see Table 1), recent studies that have used this coping style paradigm to test the relationship between behavior and HPA axis reactivity or activity in free-living animals have found that the proactive-reactive continuum is not predictive of the physiological stress response (Garamszegi et al. 2012; Ferrari et al. 2013; Dosmann et al. 2015). For example, though laboratory selection line results are consistent with predictions of the unidimensional model, when testing Great Tits in the laboratory with natural, non-selected variation in exploratory behavior, the relationship no longer holds (Baugh et al. 2012).

**Table 1.** Review of field studies testing for the covariance in behavioral coping styles and HPA activity. This non-comprehensive table includes studies conducted with natural populations testing the unidimensional (Koolhaas et al. 1999b) and two-tier models (Koolhaas et al. 2010) of the among individual relationship between behavioral coping styles (‘Behavioral Trait’) and HPA axis activity (‘Physiological Measurement’). The first section includes studies that support the main prediction from the unidimensional model that more proactive individuals have lower HPA axis activity. The second section includes studies that are contrary to the main prediction from the unidimensional model that more proactive individuals would have lower HPA axis activity. These studies do show that behavior and HPA activity covary, but the relationship is in the opposite of the direction predicted by the unidimensional model with proactive individuals having lower HPA axis activity. The third section includes studies that support the two-tier model that predicts that behavior and HPA axis activity do not covary in either direction. Correlations or estimates are included if available in the corresponding manuscript. Confidence/credible intervals are included if available; if not, p-values are included when available. If the confidence/credible interval overlapped zero, we interpreted this as the behavior trait measured and the physiological measurement did not covary. Non-true baseline samples were the first sample collected but involved some handling or trapping stress. (n = sample size of individuals, NS = non-significant (*p* > 0.05), FCM = fecal cortisol metabolites, DEX = dexamethasone (synthetic corticosteroid that exerts negative feedback on the HPA axis), ACTH = adrenocorticotropic hormone that increases adrenocortical activity)

| Species | Behavioral Trait | Physiological Measurement | n | Correlation or Estimate [CI] |
| --- | --- | --- | --- | --- |
| ***Evidence supporting the predictions of the unidimensional model that behavioral and physiological traits are negatively correlated*** |  |  |  |  |
| Great Tits (*Parus major*)<br>(Baugh et al. 2017) | Exploration | Blood corticosterone (*ACTH challenge induced*) | 85 | $R^2 = 0.051, p = 0.04$ |
| Great Tits (*Parus major*)<br>(Baugh, van Oers, Naguib, & Hau, 2013) | Exploration | Blood corticosterone (*90-min handling-restraint stress-induced*) | 16 | $b = -0.417, p = 0.015$ |
| | Exploration | Blood corticosterone (*area under the stress-induced curve*) | 16 | $b = -24.66, p = 0.007$ |
| Belding's ground squirrels (*Urocitellus belingi*)<br>(Clary et al. 2014) | Vigilance | FCM (*nominal baseline*) | 12 | $b = 2.109 [0.17, 4.05]$ |
| Brook charr (*Salvelinus fontinalis*)<br>(Farwell et al. 2014) | Activity | Whole-body cortisol (*baseline and handling/novel object stress-induced samples*) | 66 | NA* |
| House Sparrows (*Passer domesticus*)<br>(Lendvai et al. 2011) | Hovering | Blood corticosterone (*30 min handling-restraint stress-induced*) | 18 | Pearson's $r = -0.58, p = 0.017$ |
| Eastern chipmunks (*Tamias striatus*)<br>(Montiglio et al. 2012) | Exploration | FCM (*coefficient of variation*) | 58 | $b = -13.68 [-27.62, -4.45]$ |
| Plateau pika (*Ochotona curzoniae*)<br>(Qu et al. 2018) | Shyness | Blood cortisol (*true baseline*) | 292 | posterior $R = 0.45 [0.09, 0.66]$ |
| ***Evidence in the opposite direction of predictions of the unidimensional model*** |  |  |  |  |
| Poeciliid fish (*Brachyrhaphis episcopi*)<br>(Archard et al. 2012) | Exploration, Activity | Water-borne cortisol (*open-field stress-induced*) | 96 | Pearson's $r = -0.29, p = 0.005$ |
| Great Tits (*Parus major*)<br>(Baugh et al. 2013) | Exploration | Blood corticosterone (*true baseline*) | 82 | $b = 0.536, p = 0.003$ |
| Alpine marmots (*Marmota marmota*)<br>(Costantini et al. 2012) | Exploration, Activity | Blood cortisol (*non-true baseline*) | 28 | $b = 0.54$,<br>Fischer *C*-statistic model selection |
| Graylag Geese (*Anser anser*)<br>(Kralj-Fišer et al. 2009) | Aggression | FCM (*handling stress-induced*) | 10 | Spearman's $r = 0.782, p = 0.008$ |
| ***Evidence supporting the predictions of the two-tier model that behavioral and physiological traits are not significantly correlated*** |  |  |  |  |
| Great Tits (*Parus major*) | Exploration | Blood corticosterone (*handling-restrain stress-induced*) | 85 | $R^2 = 0.03, p = 0.14$ |
| (Baugh, Davidson, Hau, & van Oers, 2017) | Exploration | Blood corticosterone (*DEX challenge response*) | 85 | $R^2 = 0.02, p = 0.20$ |
| Belding's ground squirrels (*Urocitellus belingi*) | Exploration | FCM (*non-true baseline*) | 12 | $b = -0.76 [-1.12, 0.98]$ |
| (Clary et al., 2014) |  |  |  |  |
| Alpine marmots (*Marmota marmota*) | Exploration, Activity | Blood cortisol (*pre-restraint to post-restraint response*) | 28 | NA** |
| (Costantini et al., 2012) |  |  |  |  |
| Belding's ground squirrels (*Urocitellus belingi*) | Activity | FCM (*non-true baseline*) | 157 | posterior R = -0.007, [-0.123, 0.095] |
| (Dosmann et al. 2015) | Exploration | FCM (*non-true baseline*) | 157 | posterior R = -0.046, [-0.175, 0.036] |
|  | Docility | FCM (*non-true baseline*) | 157 | posterior R = 0.016, [-0.077, 0.139] |
| Alpine marmots (*Marmota marmota*) | Activity | Blood cortisol (*non-true baseline*) | 146 | posterior R = 0.04, [-0.56, 0.71] |
| (Ferrari et al. 2013) | Impulsivity | Blood cortisol (*non-true baseline*) | 146 | posterior R = 0.08, [-0.68, 0.62] |
|  | Docility | Blood cortisol (*non-true baseline*) | 146 | posterior R = 0.14, [-0.64, 0.63] |
| Collard Flycatchers (*Ficedula albicollis*) | Novel object avoidance | FCM (*non-true baseline*) | 51 | Pearson's r = -0.017, [-0.291, 0.260] |
| (Garamszegi et al. 2012) | Aggression | FCM (*non-true baseline*) | 56 | Pearson's r = -0.076, [-0.332, 0.191] |
|  | Risk-taking | FCM (*non-true baseline*) | 54 | Pearson's r = 0.074, [-0.198, 0.335] |
| Nazca Boobies (*Sula granti*) | Gardening (non-social) | Blood corticosterone (*true baseline*) | 222 | $b = 0.26, [-0.09, 0.61]$ |
| (Grace and Anderson 2014) | Shaking (non-social) | Blood corticosterone (*true baseline*) | 222 | $b = -0.06, [-0.25, 0.13]$ |
| | Aggression (non-social) | Blood corticosterone (*true baseline*) | 222 | $b = 0.19, [-0.07, 0.45]$ |
| | Gardening (non-social) | Blood corticosterone (*maximum value across 4 time points during capture-restraint*) | 222 | $b = -0.30, [-0.98, 0.38]$ |
| | Shaking (non-social) | Blood corticosterone (*maximum value across 4 time points during capture-restraint*) | 222 | $b = -0.08, [-0.44, 0.28]$ |
| | Aggression (non-social) | Blood corticosterone (*maximum value across 4 time points during capture-restraint*) | 222 | $b = -0.07, [-0.35, 0.21]$ |
| | Gardening (non-social) | Blood corticosterone (*area under the curve across 4 time points during capture-restraint*) | 222 | $b = -0.16, [-0.7, 0.38]$ |
|  | Shaking (non-social) | Blood corticosterone (*area under the curve across 4 time points during capture-restraint*) | 222 | NA, CI in figure includes 0 |
|  | Aggression (non-social) | Blood corticosterone (*area under the curve across 4 time points during capture-restraint*) | 222 | NA, CI in figure includes 0 |
| | Gardening (social) | Blood corticosterone (*true baseline*) | 222 | $b = 0.20, [0.02, 0.38]$ |
| | Shaking (social) | Blood corticosterone (*true baseline*) | 222 | $b = -0.04 [-0.18, 0.1]$ |
|  | Aggression (social) | Blood corticosterone (*true baseline*) | 222 | b = -0.02 [-0.11, 0.07] |
|  | Gardening (social) | Blood corticosterone (*maximum value across 4 time points during capture-restraint*) | 222 | b = -0.03 [-0.23, 0.17] |
|  | Shaking (social) | Blood corticosterone (*maximum value across 4 time points during capture-restraint*) | 222 | b = 0.04 [-0.2, 0.28] |
|  | Aggression (social) | Blood corticosterone (*maximum value across 4 time points during capture-restraint*) | 222 | b = -0.01 [-0.02, 0] |
|  | Gardening (social) | Blood corticosterone (*area under the curve across 4 time points during capture-restraint*) | 222 | b = 0.02 [-0.15, 0.19] |
|  | Shaking (social) | Blood corticosterone (*area under the curve across 4 time points during capture-restraint*) | 222 | NA, CI in figure includes 0 |
|  | Aggression (social) | Blood corticosterone (*area under the curve across 4 time points during capture-restraint*) | 222 | NA, CI in figure includes 0 |
| Graylag Geese (*Anser anser*)<br>(Kralj-Fišer et al. 2009) | Aggression | FCM (*non-true baseline*) | 10 | Spearman's r = -0.503, NS |
|  | Sociability | FCM (*non-true baseline*) | 10 | Spearman's r = -0.212, NS |
|  | Sociability | FCM (*handling stress-induced*) | 10 | Spearman's r = -0.127, NS |
| House Sparrows (*Passer domesticus*)<br>(Lendvai et al. 2011) | Exploration | Blood corticosterone (*30 min handling-restraint stress-induced*) | 18 | Pearson's r = -0.07, *p* = 0.797 |
|  | Shy-bold | Blood corticosterone (*30 min handling-restraint stress-induced*) | 18 | Pearson's r = -0.04, *p* = 0.866 |
|  | Exploration | Blood corticosterone (*true baseline*) | 18 | Pearson's r = -0.18, *p* = 0.487 |
|  | Shy-bold | Blood corticosterone (*true baseline*) | 18 | Pearson's r = -0.07, *p* = 0.801 |
|  | Hovering | Blood corticosterone (*true baseline*) | 18 | Pearson's r = -0.14, *p* = 0.573 |
| Plateau pika (*Ochotona curzoniae*)<br>(Qu et al. 2018) | Docility, in cage | Blood cortisol (*true baseline*) | 292 | posterior R = 0.25 [-0.02, 0.63] |
|  | Docility, in bag | Blood cortisol (*true baseline*) | 292 | posterior R = -0.06 [-0.28, 0.31] |
|  | Exploration | Blood cortisol (*true baseline*) | 292 | posterior R = -0.16 [-0.52, 0.05] |
|  | Docility, in cage | Blood cortisol (*40 min post-capture change from baseline*) | 292 | posterior R = 0.22 [-0.06, 0.47] |
|  | Docility, in bag | Blood cortisol (*40 min post-capture change from baseline*) | 292 | posterior R = 0.16 [-0.07, 0.48] |
|  | Exploration | Blood cortisol (*40 min post-capture change from baseline*) | 292 | posterior R = 0.10 [-0.42, 0.24] |
|  | Shyness | Blood cortisol (*40 min post-capture change from baseline*) | 292 | posterior R = -0.23 [-0.28, 0.22] |

\* Used AICc model selection approach. All top models included activity as predictor of cortisol. Based on figure in text, the top model indicates more active individuals have less whole-body cortisol. \*\* Used nested path model selection approach. Based on figure in text, the top model does not include a relationship between change in cortisol and coping style.

Discrepancies in the lab between recent observed relationships (Koolhaas et al. 2007) and the simple unidimensional model (Koolhaas et al. 1999), have recently led to the development of a ‘two-tier’ coping style model. This two-tier model proposes that individuals in a population can vary independently in both behavioral responses and physiological responses to environmental challenges (Koolhaas et al. 2010). This model of coping styles reframed the original model to establish behavioral coping strategies on a continuum independent of physiological coping strategies (Koolhaas et al. 2010). The distinction between the unidimensional and two-tier coping style models is significant in assessing the ecological and evolutionary consequences of variation in response to stressors. If the phenotypic correlation between behavioral and physiological stress responses (assumed by the unidimensional coping styles model) reflects an underlying correlation mediated by the effects of a hormone, this may present a limitation in the ability of populations to adapt to changing environmental stressors (Sih et al. 2004; Dantzer and Swanson 2017). Alternatively, if the two-tier model of coping style is supported, and there are two separate axes for behavioral and physiological stress responses, this suggests the potential for each trait to be an independent target of selection, potentially facilitating rapid adaptation to new environmental challenges (McGlothlin and Ketterson 2008; Ketterson et al. 2009). Exploring how coping styles relate to the physiological stress response in wild populations allows us to test across the entire spectrum of naturally occurring individual variation in behavioral coping styles, thus informing our perspective on how these mechanisms function in wild populations (Réale et al. 2007; Ferrari et al. 2013).

We investigated the relationship between three behavioral traits and one measure of HPA axis activity (concentrations of fecal cortisol metabolites, FCM) in a natural population of North American red squirrels (*Tamiasciurus hudsonicus*, hereafter, ‘red squirrels’). Previous studies in this species showed that there was a repeatable, correlated suite of behavioral traits, specifically aggression, activity, and docility, across the adulthood of an individual (Boon et al. 2007; Taylor et al. 2012). These suites of behavioral traits can also be placed along the proactive-reactive continuum as coping styles, with the more active, aggressive, and less docile individuals at the proactive end of the continuum. Differences in coping styles in red squirrels have clear environment-dependent fitness correlates (Boon et al. 2007, 2008; Taylor et al. 2014), and variation in heritable coping styles among individuals (Taylor et al. 2012) in this population may be maintained through fluctuating selection caused by changing environmental conditions (Taylor et al. 2014).

We used fecal samples as a non-invasive proxy for HPA axis activity and reactivity, which is unaffected by trapping-induced stress (Dantzer et al. 2010). In red squirrels, FCM is representative of the circulating plasma cortisol over the past ∼12 hours, with a 10.9 ± 2.3 hours lag time to peak excretion following experimental administration of cortisol (Dantzer et al. 2010). Influences of the circadian rhythm on circulating cortisol are not detected in fecal samples collected throughout the day (Dantzer et al. 2010). Additionally, glucocorticoid concentrations in fecal samples have been shown to be representative of HPA activity and reactivity (Sheriff et al. 2011; Palme 2019).

It is important to note that glucocorticoids are metabolic hormones and only one mediator of the reactive physiological stress response of an individual (Romero et al. 2009). However evidence for both the unidimensional and two-tier models of coping styles specifically connect glucocorticoids with the behavioral response (Table 1), in addition to catecholamines (reviewed in Koolhaas et al. 1999, 2010). While this is not a perfect measure of the overall physiological stress response of an individual, glucocorticoids are an important physiological mediator of the multifaceted stress response (Sapolsky et al. 2000; Romero et al. 2009). Glucocorticoids are secreted to mobilize energy in response to a stressor in the environment, but also exert pleiotropic effects (Sapolsky et al. 2000). For example, fluctuating baseline glucocorticoids act as a mediator of future reproductive investment in European Starlings (*Sturnus vulgaris*) by preparing individuals for energetically expensive reproductive seasons (Love et al. 2014).

To test the unidimensional and two-tier models of the overall stress response, we measured FCM as a non-invasive marker of HPA axis activity and the behavior of individuals using three behavioral assays (open-field trial, mirror-image stimulation trial, and handling docility assay) to measure coping style. We then compared the FCM concentrations to the behavioral coping style of individual squirrels. A relationship between FCMs and behavioral coping style across individuals would support the unidimensional model, whereas a lack of relationship would support the two-tier model.

## Methods

### Study species

North American red squirrels are a sexually monomorphic species of arboreal squirrels (Boutin and Larsen 1993). Females and males are both territorial of their food-cache (located on the center of their territory) year-round (Dantzer et al. 2012; Siracusa et al. 2017). Red squirrels in the region of our study rely on seeds produced by white spruce (*Picea glauca*) trees as their primary food source (Fletcher et al. 2010). Squirrel population density is closely associated with mast seeding of the white spruce, or episodes of booms and busts in food availability (McAdam and Boutin 2003; Fletcher et al. 2010; Dantzer et al., 2013). Red squirrels have one litter per year, with the exception of mast years when autumn spruce seed is superabundant (Boutin et al. 2006; McAdam et al. 2007).

Our study was conducted as a part of the Kluane Red Squirrel Project, a long-term study of wild population of red squirrels within Champagne and Aishihik First Nation traditional territory along the Alaska Highway in the southwest Yukon, Canada (61**°**N, 138**°**W). Each squirrel was tagged with a unique set of alphanumeric stamped ear tags (National Band and Tag Company, Newport, KY, USA). At each live-trapping (Tomahawk Live Trap, Tomahawk, WI, USA) event, body mass and reproductive status of the squirrel were recorded. Female reproductive status was determined through changes in body mass, by nipple condition, and by abdominal palpations of developing fetuses in females. Male reproductive status was determined by palpating for the presence of testicles either in the scrotum (breeding) or abdomen (non-breeding). A more detailed description of the population and general methods can be found in McAdam et al. (2007).

The local population of red squirrels was broken down into three study populations in different locations: two were control populations (referred to hereafter as ‘control grids’) and one was provided with supplemental food between 2004 and 2017, such that squirrel density was increased (Dantzer et al., 2013; hereafter referred to as ‘high-density grid’). Squirrels on the high-density grid were provided with 1 kg of peanut butter (no sugar or salt added) approximately every six weeks from October to May (Dantzer et al. 2012). We included these squirrels to increase our sample size, and included study grid type as a covariate in all models to control for variation between the grids. Additionally, high conspecific competition is a significant environmental factor influencing the physiological stress response of red squirrels (Dantzer et al. 2013) and so was important to include as a covariate in our statistical models (Table 2). All work was conducted under the animal ethics approvals from Michigan State University (AUF#04/08-046-00) and University of Guelph (AUP#09R006).

**Table 2.**
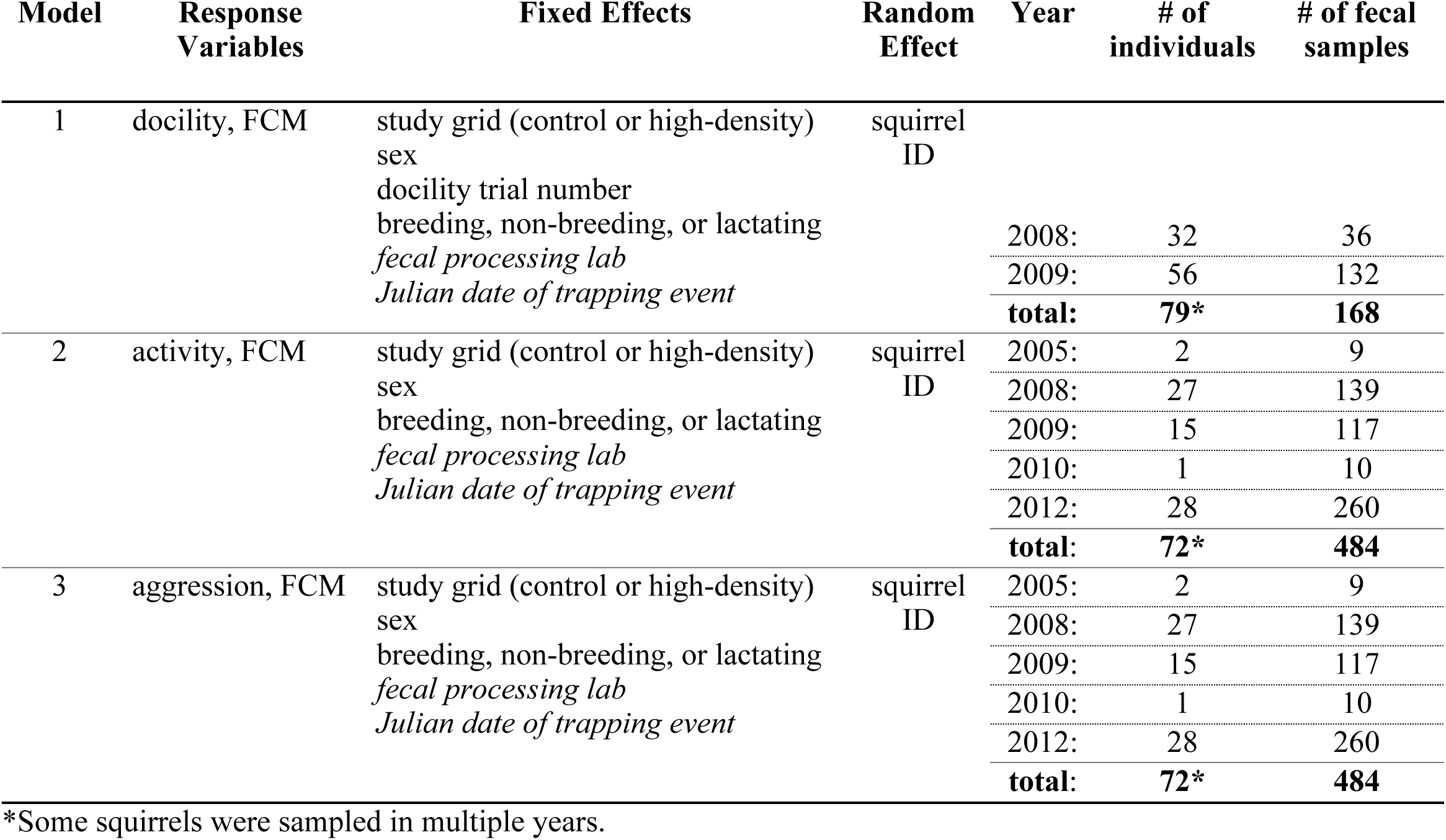
Bivariate model structures and sample sizes. This table breaks down the sample sizes and the variables (fixed and random effects) considered in each bivariate mixed effects model. Response variables shown are for the bivariate models that tested the association between behavioral traits (docility, activity, or aggression) and a measure of the physiological stress response (fecal cortisol metabolites or FCM). Italics indicate fixed effects estimated for FCM concentration only.

### Behavioral trials

Squirrels were subjected to two behavioral trials to measure ‘activity’ and ‘aggression’: an open-field (OF) trial, and a mirror image stimulation (MIS) trial (Boon et al. 2007; Taylor et al. 2012). These behavioral trials were conducted for other projects and were not evenly distributed across years. We performed trials in 2005, 2008, 2009, 2010, and 2012 (see Table 2 for a breakdown of sample sizes). During these years, additional trials were conducted on squirrels in this population for multiple studies (see Boon et al. 2007; Boon et al. 2008; Kelley et al. 2015; Taylor et al. 2012, 2014), but for the purposes of this analyses, we only included trials for which we also had FCM concentration data for that individual. All squirrels were mature adults (>1 year old) at the time of the trial.

To measure an individual’s coping style, we conducted OF and MIS trials during the same trapping event, with the OF trial completed first followed by the MIS trial. Squirrels habituate to these tests (Archer 1973; Boon et al. 2008; Martin and Réale 2008), but the behavior of individual squirrels over time is known to be repeatable (Boon et al. 2007, 2008; Taylor et al. 2014). For simplicity, we thus used only the results of each individual’s first test as a measure of its activity and aggression. We transferred focal squirrels from a live trap into the arena using a canvas handling bag. The same portable testing arena was used for both trials, and consisted of a 60 × 80 × 50 cm white corrugated plastic box with a clear acrylic lid (Taylor 2012). Four blind holes made with black PVC caps in the bottom of arena allowed the squirrel to explore possible ‘escape routes’. We exposed a 45 × 30 cm mirror fixed to one end of the arena after the OF trial to begin the MIS trial. A digital video camera recorded behavior in the arena. We performed all behavioral trials on the territory of the focal individual. Between trials, we cleaned the arena using 70% isopropyl alcohol.

To quantify behavior from the videos, we used manual scoring methods with an ethogram developed and used in previous red squirrel studies (Boon et al. 2007, 2008; Taylor et al. 2012, 2014; Kelley et al. 2015; Supplementary Material Table S1). Because these videos were collected and scored across multiple years, observers used different software programs depending on what program was available the year they were scored. Trials conducted in 2005 were scored using The Observer Video-Pro 5.0 (Noldus Information Technology, Wageningen, The Netherlands). Trials conducted from 2008-2010 were scored using JWatcher Video 1.0 (Blumstein and Daniel 2007). Trials conducted in 2012 were scored using Cowlog software (Hänninen and Pastell 2009). Regardless of the software used, the ethogram and the overall method of scoring the videos remained consistent. Because this is a manual process and the software simply records keystrokes indicating behaviors observed, it is not likely that the software used impacted the score. Furthermore, a previous study using some of our dataset showed high inter-observer reliability for the behavioral measures we recorded from these videos (Taylor et al., 2012), so it again seems unlikely that the software used would influence the behavioral data we extracted from the videos. During the OF trial, we recorded the mutually exclusive behaviors of time spent walking, sniffing, chewing, rearing, grooming, and being still. Additionally, we recorded the number of jumps and head-dips in the false holes. During the MIS trial, we recorded the amount of time spent in the third of the arena closest to the mirror, and the amount of time spent in the third of the arena farthest from the mirror. We also recorded the number of aggressive contacts with the mirror (attacks), the latency until the first attack, and the latency until the first approach towards the mirror. A detailed description of the video scoring methods can be found in Boon et al. (2008). Following Taylor et al. (2012), behaviors with an inter-observer reliability of greater than 0.7 were used in analyses (see Supplementary Material Table S1 and S2 for a list of behaviors used in the analyses).

As an additional behavioral measurement, we also measured ‘docility’ of individual squirrels. In 2008 and 2009, docility measurements were collected on many squirrels for other studies (see Boon et al. 2007; Taylor et al. 2012), but for the purposes of this study, we focused only on trials that were conducted during a trapping event where fecal samples were also collected and subsequently analyzed (n = 168 trapping events). We quantified docility as the squirrel’s response to handling (for examples in other species, see Carere and Oers 2004; Martin and Réale 2008; Montiglio et al. 2012). We transferred squirrels from the trap into a canvas handling bag and placed the squirrel on a flat surface. We measured docility during handling by counting the number of seconds out of 30 seconds in which the squirrel was not struggling. A squirrel that spent most of the time immobile during the test was considered docile, a trait previously demonstrated to be repeatable (Boon et al. 2007; Taylor et al., 2012) and heritable (Taylor et al., 2012) in this population. This test was conducted an average of 8 (min = 1, max = 42) times on 79 individual squirrels caught between 2008 and 2009. Docility scores were z-scored for analyses. See Table 2 for detailed sample sizes.

### Fecal cortisol metabolites

From 2005 to 2014, we opportunistically collected a total of 703 fecal samples during routine trapping of squirrels with peanut butter for measurement of FCM concentrations corresponding to individuals with behavioral data (see Dantzer et al. 2010). Fecal samples were collected from under live-traps within two hours of trapping and placed in 1.5 mL vials stored in a −20 °C freezer within five hours of collection. Urine contaminated feces were excluded. All fecal samples were lyophilized for 14-16 h before being pulverized in liquid nitrogen using a mortar and pestle. Using 0.05 g of dry ground feces, steroid metabolites were extracted by adding 1 mL of 80% methanol and vortexing samples at 1450 RPM for 30 min, and then centrifuging for 15 min at 2500 g (Dantzer et al. 2010; Palme et al. 2013). The resulting supernatant was stored at −20 °C for analysis via glucocorticoid metabolite assay using a 5α-pregnane-3β,11β,21-triol-20-one antibody enzyme immunoassay (EIA; see Touma et al. 2003). A detailed validation and description of steroid extraction and EIA with red squirrel fecal samples can be found in Dantzer et al. (2010).

Fecal samples were analyzed across multiple assays and in two different labs (n = 355 at University of Toronto Scarborough and n = 348 at University of Michigan) but using the same protocol. We confirmed that our measures of FCM concentrations were highly repeatable across assays or labs through the following. First, a separate group of fecal samples (n = 128 samples) were analyzed in both labs and the optical density of these samples were closely correlated (Pearson correlation = 0.88). This indicates that the data were comparable, but we also included a covariate in our statistical models for where the data were analyzed (see below). Second, using pooled samples that were run repeatedly on different plates (n = 115), we found that the estimates of optical density for these pool samples were highly repeatable (*R* = 0.85, 95% CI = 0.54-0.93). Finally, using a linear mixed-effects model, we partitioned the variance in the optical density recorded for the pooled samples that were run across these different plates. We found that most of the variance was due to the sample itself (85.1%) with relatively little of it being explained by intra-assay variation as all samples were run in duplicate (4.9%) or by inter-assay variation (9.9%). Together, this indicates that our measures of FCM concentrations should be comparable across assays and across labs. See Table 2 for a representation of how sample sizes were broken down in each dataset.

### Statistical methods

All statistical analyses were conducted in R version 3.4.3 (R Core Team 2016). For the OF and MIS trials, we used two principal components analyses to reduce the redundancy among behavioral measurements and calculate composite behavioral scores for each trial, as we have done previously in this system (Boon et al. 2007, 2008; Taylor et al. 2012, 2014; Kelley et al. 2015). To conduct the principal components analyses with correlation matrices, we used the R package ‘ade4’ version 1.7-10 (Dray and Dufour 2007). By reducing the multiple behaviors observed down to one metric for each trial, we were able to assess the primary variation among individuals along those axes. All further analyses used the scores calculated from the principal component loadings (Supplementary Material Table S2) for each trial. From the OF trial, we interpreted the first principal component as a measure of overall ‘activity’, as it has previously been interpreted in this population (Boon et al. 2007, 2008; Taylor et al. 2012, 2014). In our data set, the first component explained 64% of the variation in behavior across OF trials. From the MIS trial, we interpreted the first principal component as a measure of ‘aggression’, as it has also previously been interpreted (Boon et al. 2007, 2008; Taylor et al. 2012, 2014). In our data set, the first component explained 60% of the variation in behavior across MIS trials.

Because we were interested in an estimate of the covariance of FCM concentrations and personality, we used a multivariate framework to conservatively address how the two types of stress responses interact, as explained in Houslay and Wilson (2017). In this study, we were interested in how FCM concentrations and behavioral traits co-varied among individuals. To investigate this in a multivariate framework, we used a Bayesian generalized linear mixed effects multivariate model based on a Markov chain Monte Carlo algorithm with the R package ‘MCMCglmm’ version 2.25 (Hadfield 2010) to assess the relationship between FCM concentrations and behavior. All fecal cortisol metabolite concentration data were ln-transformed to improve normality of residuals.

For docility analyses, we used measurements of docility paired with the FCM concentrations of that trapping event. Using a bivariate generalized linear mixed-effects model, we asked whether individuals with higher mean FCM concentrations have higher mean docility scores (among-individual covariance), and whether individual observations of FCM concentration and docility relative to the individual’s mean concentrations were correlated (within-individual covariance). Within-individual covariance indicates how the FCM concentrations and docility scores of one individual covary across multiple observations for that individual; in essence if we have multiple unique measurements of FCM concentration and docility from one individual, does the docility score predict FCM concentrations on that day? In contrast, among-individual covariance measures the relationship between FCM concentration and docility across individuals in the population; in other words, does an individual’s average docility score predict its average FCM concentration? The bivariate model for docility and FCM concentration included fixed effects of study grid (control or high-density), sex, Julian date of trapping event (continuous), trial number, breeding status (breeding/non-breeding/lactating), and a variable to indicate where the fecal sample was processed (UT Scarborough/UM). Docility measurements were taken across multiple trapping events for a squirrel, therefore we included trial number to control for any variation caused by habituation to the process of being trapped and restrained in the bag (Boon et al., 2007; Taylor et al., 2012). We specified in the model to estimate the fixed effects of Julian date of trapping event and the location of fecal sample processing for only FCM concentration. These fixed effects were included because they have previously been shown to influence FCM concentration in red squirrels (Dantzer et al. 2010, 2013), and thus were included to control for variation among these variables. Although the correlation between UT Scarborough and UM samples was high (0.88), we included location of fecal sample processing to account for any minor variation between the locations.

Because we only had one measurement of aggression and activity per individual, we were unable to estimate the within-individual covariance between FCM concentration and the activity/aggression of that individual. Thus, the bivariate models for activity and aggression were structured to control for only one individual activity and aggression score per individual, and only estimate among-individual covariance. The models for activity and aggression included the same fixed effects as the model for docility, except trial number was not included. All trials were the first trial in the lifetime of that individual, so there is no variance in novelty of the arena across individuals. Again, we estimated fixed effects of the Julian date of trapping event and location of fecal sample processing for only FCM concentration. With this model structure, we were able to more precisely control for variation in FCM concentration due to reproductive condition and time of year.

We fit all bivariate MCMCglmm models with uninformative priors (as in Houslay and Wilson 2017) for 2,100,000 iterations with the first 100,000 discarded, and 1 out of every 1,000 of the remaining iterations used for parameter estimations. Credible intervals (95%) around the correlation were based on the MCMC chain iterations. To confirm convergence using a combination of methods, as suggested in a comparative review (Cowles and Carlin 1996), we ran all MCMCglmm models three times for comparison using the Gelman-Rubin statistic (Gelman and Rubin 1992), and we also ran the Geweke diagnostic (Geweke 1992). All models passed both diagnostics for convergence.

### Data availability

The datasets analyzed during the current study available from the corresponding author on reasonable request.

## Results

Our results indicate that docility, activity, and aggression did not co-vary with FCM concentrations among individuals. Using a bivariate generalized linear mixed effects model approach, the within-individual covariance indicated that an individual’s FCM concentrations did not correlate with docility (*r* = 0.020, CI = [−0.14, 0.28]). Our models also indicated that among individuals, FCM concentrations did not correlate with docility (*r* = 0.14, CI = [−0.64, 0.83], Figure 1), activity (*r* = 0.15, CI = [−0.17, 0.47], Figure 2) or aggression (*r* = 0.29, CI = [−0.098, 0.56], Figure 2) (Table 3). Regardless of the statistical significance of these relationships, the direction of the observed effect was opposite to the predicted relationship between behavior and FCM concentrations. The direction of these correlations suggest more active and more aggressive squirrels may have higher FCM concentrations, but this is not conclusive due to confidence intervals overlapping zero.

**Figure 1.**
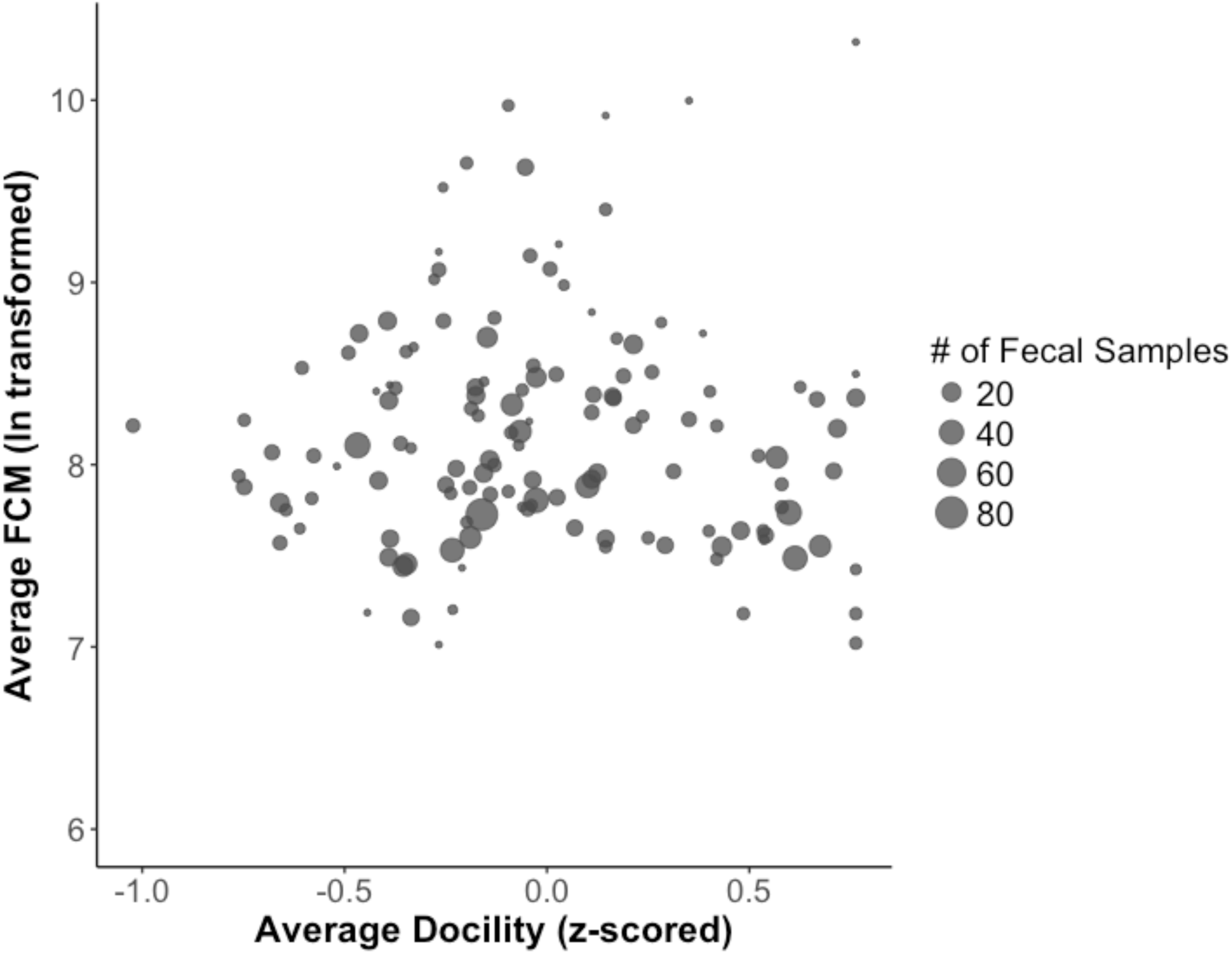
Stress reactivity and activity, as measured by average FCM concentration, is not predicted by docility in North American red squirrels. Size of the points represents number of fecal samples included for that individual.

**Figure 2.**
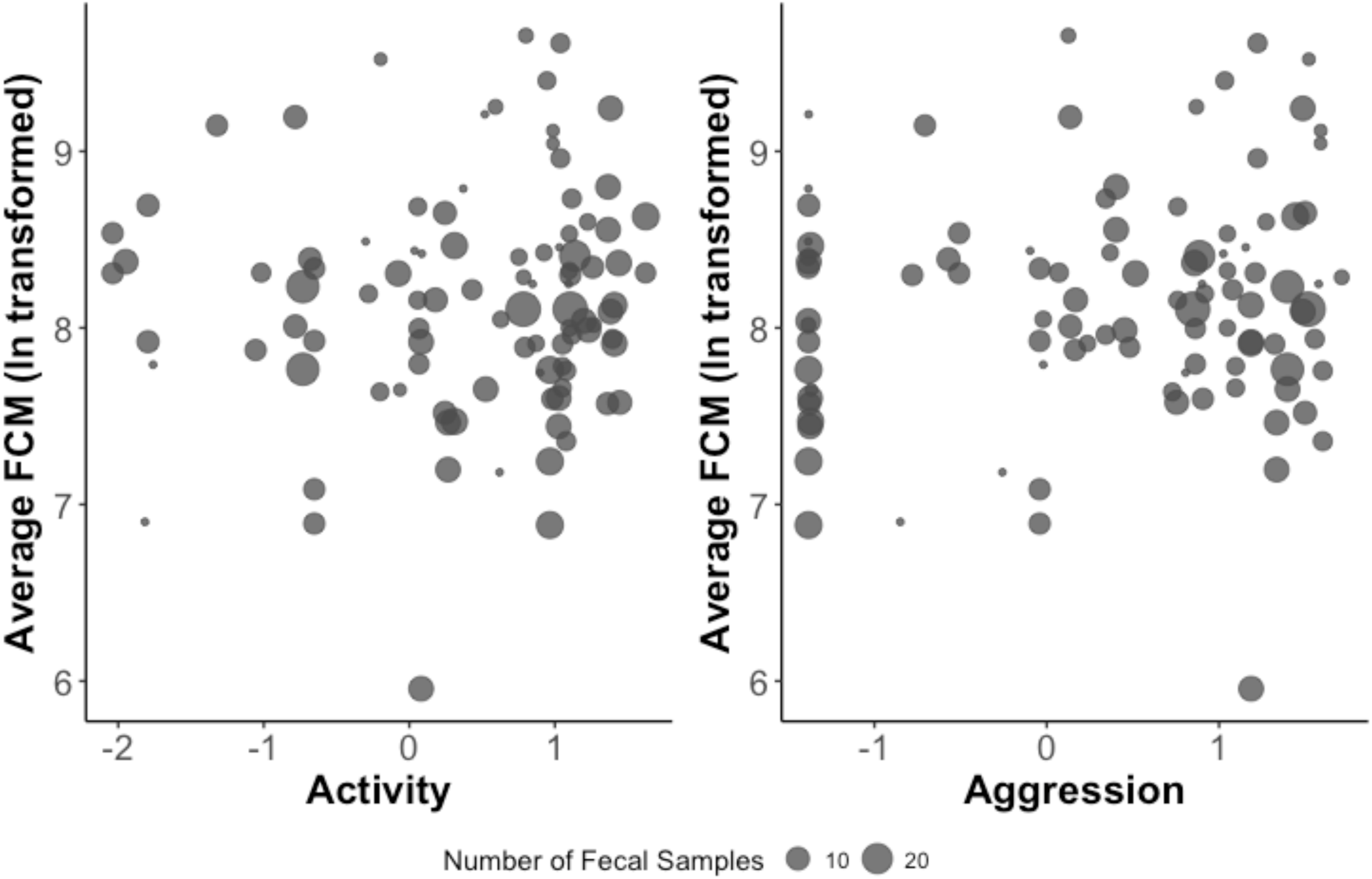
Stress reactivity and activity, as measured by average FCM concentration, is not predicted by activity or aggression in North American red squirrels. Activity and aggression are from scores determined by the principal component analyses. Size of the points represents number of fecal samples included for that individual.

**Table 3.**
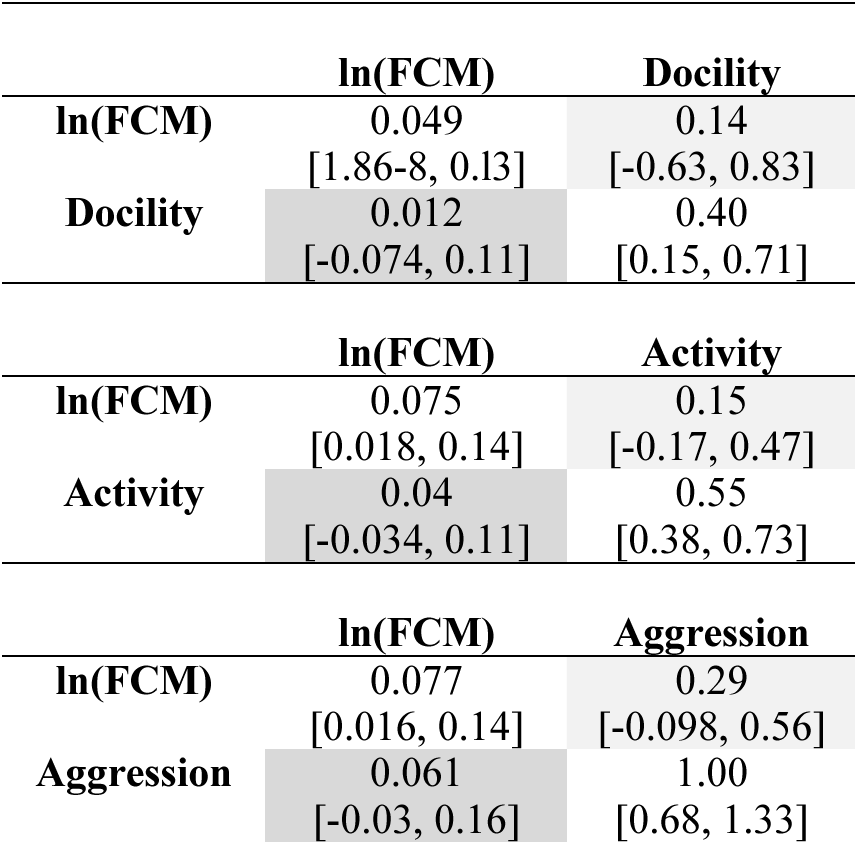
Multivariate results for relationships between FCM and behaviors. Results from our three bivariate generalized linear mixed-effects model models to examine the relationships between FCM and each of three behaviors individually (activity and aggression models: n = 484 fecal samples; docility model: n = 168 fecal samples). Among-individual variances are listed on the diagonal, covariances below and correlations above (with the lower and upper bounds of 95% CIs in parentheses).

## Discussion

We demonstrated that the behavioral coping style (represented by three behaviors) and one measure of the physiological stress response (FCM concentrations) did not co-vary in a free-ranging mammal. Independent variation between the behavioral and physiological stress responses supports the two-tier model of coping strategies proposed by Koolhaas et al. (2010). This model proposes that within a species, individuals can exhibit a consistent behavioral response anywhere along the proactive-reactive continuum but independent of their physiological stress response, which can range from a low to a high HPA axis activity. Contrary to many studies (Raulo and Dantzer, 2018), proactive, or highly active/aggressive red squirrels did not always exhibit lower HPA axis activity than reactive individuals. In fact, the parameter estimates were in the opposite direction from those predicted by the unidimensional model. Specifically, the unidimensional model predicts that a more active behavioral stress response and HPA axis activity should be negatively correlated and we found that they were instead positively correlated, though again these confidence intervals overlapped zero. Although we did find that the three behavioral measures were phenotypically correlated (see Supplementary Material), physiological stress, as measured by FCM concentration, does not appear to be the pleiotropic mechanism causing this covariation.

Previous studies that have found behavior and HPA axis reactivity are linked have used a different statistical framework than our study. Thus, it is possible our results are simply the outcome of using a more conservative statistical test. However, our results were robust across statistical techniques as we also ran the same models using a different statistical technique (linear mixed effects models) that has been used in previous studies (e.g. Lendvai et al. 2011; Montiglio et al. 2012). These results from the linear mixed effects models (Supplementary Material Table S5) and those from the bivariate models (presented above) both support the hypothesis that behavior and physiology are independent in our study.

Our study contributes to an emerging trend of a lack of a strong relationship between behavioral and physiological stress responses in wild and laboratory animals (reviewed by Raulo and Dantzer 2018). For example, wild alpine marmots (*Marmota marmota*) exhibit a lack of among-individual correlation between activity and plasma cortisol concentrations, as well as between docility and plasma cortisol (Ferrari et al. 2013). Likewise, docility and exploration were not correlated with a change in plasma cortisol in response to a stressor in plateau pika (*Ochotona curzoniae*: Qu et al. 2018). Additional studies measuring fecal glucocorticoid metabolites demonstrate that HPA axis activity does not correlate with shy-bold behavioral types in wild flycatchers (*Ficedula albicollis*; Garamszegi et al. 2012), or with exploration/activity in Belding’s ground squirrels (*Spermophilus beldingi*; Dosmann et al. 2015). In captivity, Holstein Friesian heifer calves (*Bos taurus*) HPA axis reactivity to ACTH is not correlated with their response to novelty (Van Reenen et al. 2005).

The unidimensional model posits that both HPA axis activity and reactivity should be lower in proactive animals (Koolhaas et al. 1999). However, it should be noted that measurements of fecal glucocorticoid metabolites in red squirrels may not allow for direct measurement of the reactivity of the HPA axis, which may correlate more strongly with behavioral stress responses compared to basal regulation (Baugh et al. 2012). Although a study on free-ranging eastern chipmunks (*Tamias striatus*) showed evidence supporting covariance of behavioral response and physiological stress response from fecal samples, the study used only one metric of physiological stress (coefficient of variation of fecal glucocorticoid metabolites) per individual (Montiglio et al. 2012). This statistical method was limiting in that it did not consider the uncertainty around each individual’s measure of HPA axis activity. Our research expands upon the chipmunk study by using more conservative statistical methods, which were not widely used until recently (Houslay and Wilson 2017), in addition to linear models and multiple behavioral assays to establish coping styles. Using both of these approaches, we showed that the behavioral coping style (comprised of three correlated behaviors) does not covary with one measure of physiological stress. We acknowledge that the studies included in Table 1 are across multiple taxa, behaviors, and HPA axis activity measurements. This likely contributes to the equivocal nature of support for these models in wild animals. Though we focused only on wild animals in our brief review (Table 1), empirical studies using laboratory animals also include variation in measurements. Due to the large variability across studies in measurements of HPA axis activity and behavior, perhaps a less generalized model of the relationship between behavior and physiological stress may be more predictive for future studies than our current models.

Our study was conducted using adult red squirrels. In this population, high juvenile mortality results in a high opportunity for selection during the first year of life (McAdam et al. 2007). Due to these strong selective pressures, we must consider the possibility that selection may have already eroded the (co)variance of physiological and behavioral stress responses in surviving adults. For example, perhaps juveniles with high covariance of the physiological and behavioral stress response were unable to adaptively respond to environmental conditions, whereas juveniles with low covariance were able to adaptively respond to conditions with the two stress responses decoupled. Additionally, these selective pressures fluctuate across years because red squirrels rely on a masting food source (white spruce) that goes through episodes of booms and busts in production of reproductive cones (McAdam and Boutin 2003; Fletcher et al. 2010). Following the masting of spruce trees, squirrel populations increase in density, which may generate density-dependent selection on juvenile traits (Dantzer et al. 2013; Fisher et al. 2017). These fluctuations in selection may maintain genetic variation in behavioral traits in this population (Taylor et al. 2014), and, if a pleiotropic hormone was the mechanism underlying these behavioral correlations, it could limit an adaptive behavioral response to this fluctuating selection if the selective forces on hormone levels and the behavior push in opposite directions (Ketterson and Nolan Jr. 1999; McGlothlin and Ketterson 2008). For example, if a high activity is beneficial in one environmental condition but high HPA axis reactivity is not, a strong correlation between the two traits would constrain an individual’s ability to show an adaptive behavioral response to the current environmental conditions. Recent work, however, suggests a hormonal pleiotropic relationship is likely not powerful enough to constrain independent evolution of two traits (Dantzer and Swanson 2017). Alternatively, if the hormone does not show a pleiotropic relationship with behavior and selection for both is working in the same direction, then the absence of a correlation could slow their adaptive response relative to a situation with a positive pleiotropic relationship between the two traits (Ketterson et al. 2009).

We also must consider the possibility that different behavioral traits are favored at different life stages. Additional work in this study system has shown wider variation in these behavioral traits among juvenile squirrels, with individuals at both extremes of the proactive-reactive continuum, and individuals regress to the mean as they age (Kelley et al. 2015). This is a potential limitation of our study. Studies using selection lines, and therefore individuals at extremes in behavioral response, may therefore be more appropriate for making predictions about juvenile red squirrels, though perhaps not appropriate for predictions about adults. In both Great Tits and rainbow trout (*Oncorhynchus mykiss*), studies using exploration selection lines have found evidence to support a correlation between behavioral and physiological stress responses (Øverli et al. 2007; Baugh et al. 2012). This suggests the potential to detect a relationship between behavioral and physiological stress responses in juvenile red squirrels, when individuals are more widely dispersed along the proactive-reactive continuum. Future work on the relationship between coping styles and physiological stress responses should investigate the ontogeny of the relationship, and how it may change across life stages.

Our study helps establish a foundation to use in exploring the fitness consequences of variability across two axes of the stress response, behavioral and physiological. Building upon this current work, we have an opportunity to explore the mechanisms contributing to each axis of variation independently. For instance, the maternal environment during ontogeny may influence the development of the physiological stress axis, or contribute to the behavioral coping style (reviewed in Meaney 2001). Furthermore, our study provides additional evidence supporting the lack of direct phenotypic correlation between behavioral and physiological stress responses in wild animals exhibiting natural variation in stress responses. Our study, in conjunction with previous studies on these models in wild animals (Table 1), suggests a need for a more generalizable model of the relationship between the behavioral and physiological stress responses, perhaps taking into account the environmental conditions experienced by the species. Further studies in wild animals are needed to explore the mechanisms underlying this variation along the phenotypic landscape of the stress response and the adaptive value of such variation.

Studies on the relationship between the behavioral and physiological stress phenotypes in wild animals in variable environments provide insight into the pleiotropic constraints on the evolutionary paths these populations may take. Our study contributes to a growing body of work in support of the two-tier model of coping styles and physiological stress reactivity and activity in wild and laboratory populations. Specifically, our study demonstrated that the FCM concentration of wild red squirrels is independent of an individual’s activity, aggression, and docility. Given that red squirrels in this region experience a fluctuating environment in terms of competitors (Dantzer et al. 2013), food (Boutin et al. 2006), and predators (O’Donoghue et al. 1998; Studd et al. 2014) and also fluctuating selection on behavioral traits (Boon et al., 2007; Taylor et al., 2014), having behavioral and physiological responses that are uncorrelated may be beneficial for adapting to this environmental variability. If similar results are found in other species, the lack of a phenotypic relationship between the behavioral and physiological stress responses could have important evolutionary implications, particularly for those species living in fluctuating environments.

## Acknowledgments

We thank Agnes MacDonald and her family for long-term access to her trapline, and the Champagne and Aishihik First Nations for allowing us to conduct our work within their traditional territory. We thank Adi Boon, Amanda Kelley, and Ryan W. Taylor for collecting much of the behavioral data and all the volunteers, field assistants, and graduate students for their assistance in data collection. This work was supported by American Society of Mammalogists to SEW; University of Michigan to SEW and BD; National Science Foundation (IOS-1749627) to BD; and Natural Sciences and Engineering Research Council to SB, AGM, JL, and RB. This is publication XX of the Kluane Red Squirrel Project.

## Ethical approval

All applicable international, national, and/or institutional guidelines for the care and use of animals were followed. All procedures performed in studies involving animals were in accordance with the ethical standards of the institution or practice at which the studies were conducted.

