## Supplemental for "Stress activity is not predictive of coping style in North American red squirrels"

### Electronic Supplemental Material

**Estimating repeatability of behavioral traits and phenotypic correlations among behavioral traits**

We used the R package ‘rptR’ version 0.9.21 (Stoffel et al. 2017) to estimate the within-individual repeatability of docility in a model lacking any fixed or random effects. Due to the count nature of the data, we performed repeatability estimation using a Poisson distribution generalized linear mixed-effects model with individual identity (ID) as a random intercept effect. We used parametric bootstrapping (n = 1000) to estimate the confidence interval for repeatability of docility.

A previous study on red squirrel personality found phenotypic and genetic correlations among activity, aggression, and docility, such that more active squirrels were more likely to be aggressive and less docile (Taylor et al. 2012). To confirm that these phenotypic correlations remained in our subset of the behavioral data used in this previous study, we tested for pairwise correlations between activity, aggression, and docility among individuals. To do this while preserving the multiple measurements of docility per individual, we used a Bayesian generalized linear mixed effects multivariate model based on a Markov chain Monte Carlo algorithm with the R package ‘MCMCglmm’ version 2.25 (Hadfield 2010). The MCMCglmm framework allowed us to test for an among-individual correlation between a trait with repeated measures (in this case, docility) and variables with only one observation per individual (in this case, activity and aggression) by controlling for the lack of within-individual variation of the behavioral trait with only one observation. This model was limited to data from individuals tested for all three behavioral traits (n = 23).

### Results

Docility scores were significantly repeatable within-individuals (R = 0.33, CI = [0.17, 0.49]). Estimates of the phenotypic correlations between our three behavioral traits were in the same direction as previous studies in this population, with active squirrels being more aggressive and less docile (Supplemental Material Table S3), though the credible intervals for the estimates were near zero or overlapped zero, likely due to the smaller sample size of individuals with data for all three behavioral traits (n = 23). These findings, combined with results from previous studies in red squirrels with much larger sample sizes (Boon et al. 2007; Taylor et al. 2012), support the idea that these behaviors are phenotypically correlated, and may be shaped by an underlying pleiotropic mechanism.

**Assessing the association between behavioral and physiological stress responses using linear mixed-effects models**

Previous studies that have assessed the association between behavioral and physiological stress responses did so using linear models (e.g. Baugh et al. 2013; Clary et al. 2014). The recently developed bivariate MCMCglmm model approach that we used here is arguably more conservative than this previously used approach (Houslay and Wilson, 2017). We wanted to compare results from this previously used linear model approach to the recently developed and perhaps more appropriate Bayesian statistical approach to this type of multivariate question (Houslay and Wilson, 2017). As a comparison to our bivariate MCMCglmm model approach presented in the main text of the Results, we also used R package ‘lme4’ version 1.1-15 (Bates et al. 2015) to create general linear models to test for relationships between each behavioral trait and average FCM concentrations with one measure per individual. For our linear models, we used the average of all ln-transformed FCM concentrations from an individual as a response variable. The relationship between behavior and hormones can be bidirectional and we do not propose any causality with these models. However, to keep our linear models consistent with our bivariate models with respect to fixed effects we structured the linear models with average ln(FCM concentration) as the response variable, in order to identify the relationship between FCM concentration and variation in behavior, while controlling for variation due to study area and sex. To estimate P-values, we used the R package ‘lmerTest’ version 2.0-36 (Kuznetsova et al. 2016). Separate models were used for each behavioral trait. Linear models for docility tested for the relationship between average docility across all trials of an individual and average FCM concentration of that individual. Fixed effects in these models predicting average FCM concentration included the behavioral trait of interest (docility, activity, or aggression), study area (control or high-density), and sex. Normality and homoscedasticity of residuals of all linear models were confirmed visually.

**Results**

Consistent with our bivariate model framework (results shown in main text), our linear models did not indicate a relationship between the behavioral coping style and physiological stress response. The models did not detect a significant relationship between average FCM concentrations and activity (b = 0.08, $t$ = 1.18, $p$ = 0.24), aggression (b = 0.11, $t$ = 0.1.66, $p$ = 0.099), or average docility (b = -0.0029, $t$ = -0.38 $p$ = 0.71) (Table S5).

**Table S1. Ethogram of open-field and mirror image stimulation trials**

Ethogram used to score behaviors in the open-field and mirror image stimulation trials, modified from Boon et al. (2007). All state behaviors for the open-field trial are mutually exclusive unless noted. In the mirror image stimulation trial, approaching the mirror is non-mutually exclusive with location in the arena.

| OF Behavior | Description |
|---|---|
| jump (E) | Jumping. |
| hang (S) | Hanging from the top of the arena. Behaviors like chew, or scan can be performed while hanging. |
| chew or scratch (S) | Scratching at or chewing the OF arena. |
| groom (S) | Paw or mouth grooming. |
| hole (E) | Interactions with one of the 4 blind holes. |
| still (S) | When the squirrel is still for 2 seconds. |
| walk (S) | When the squirrel is moving around the arena. |

| MIS Behavior | Description |
|---|---|
| attack mirror (E) | Each time the squirrel aggressively contacts the mirror |
| approach mirror (S) | When the squirrel moves in the direction of the mirror. |
| front (S) | When the squirrel enters the front of the arena |
| back (S) | When the squirrel enters the back of the arena |

(E) indicates an event behavior and (S) indicates a state behavior.

**Table S2. Principal component analyses loadings**

Principal component analysis results from the open-field and mirror image stimulation trials.

Loadings greater than 0.2 were used to interpret the components and are italicized.

| **OF Behavior** | **Loading** |
|---|---|
| *time spent walking* | *0.327* |
| *time spent hanging* | *0.260* |
| chewing or scratching | 0.134 |
| number of jumps | 0.128 |
| hole head dips | 0.019 |
| time spent grooming | 0.005 |
| *not moving* | *-0.888* |

| **MIS Behavior** | **Loading** |
|---|---|
| *time spent in front of arena* | *0.422* |
| number of attacks | 0.142 |
| *time spent in back of arena* | *-0.434* |
| *latency to attack* | *-0.499* |
| *latency to approach* | *-0.603* |

**Table S3. Multivariate results for relationships between all behaviors**

Results from our multivariate model using a subset of data to examine the relationships between docility, activity, and aggression (n = 23 squirrels). Among-individual variances are listed on the diagonal, covariances below and correlations above (with the lower and upper bounds of 95% CIs in brackets).

| | Docility | Activity | Aggression |
|---|---|---|---|
| **Docility** | 0.68<br>[0.30, 1.26] | -0.42<br>[-0.77, -0.006] | -0.22<br>[-0.60, 0.14] |
| **Activity** | -0.33<br>[-0.74, 0.13] | 0.91<br>[0.42, 1.47] | 0.12<br>[-0.30, 0.50] |
| **Aggression** | -0.23<br>[-0.65, 0.20] | 0.14<br>[-0.45, 0.63] | 1.41<br>[0.83, 2.16] |

### Table S5. Full results of linear models

Full results from three linear models between average FCM concentration and each behavioral trait independently. Significant fixed effects ($p < 0.05$) are in bold. Grid type compares the high-density grid to the control (intercept) grid. Sex shows how males differ from females (intercept).

| Response Variable | Fixed Effect | b | SE | t | *p*-value |
|---|---|---|---|---|---|
| **Average FCM (ln transformed)** | | | | | |
| n = 72 individuals | **Intercept** | **8.04** | **0.081** | **99.28** | **< 0.0001** |
| | Activity | 0.081 | 0.069 | 1.18 | 0.24 |
| | **Grid type** | | | | |
| | **high density** | **0.44** | **0.18** | **2.43** | **0.017** |
| | Sex | | | | |
| | males | 0.086 | 0.20 | 0.43 | 0.67 |
| **Average FCM (ln transformed)** | | | | | |
| n = 72 individuals | **Intercept** | **8.03** | **0.079** | **102.18** | **< 0.0001** |
| | Aggression | 0.11 | 0.063 | 1.66 | 0.099 |
| | **Grid type** | | | | |
| | **high density** | **0.44** | **0.18** | **2.50** | **0.014** |
| | Sex | | | | |
| | males | 0.14 | 0.20 | 0.70 | 0.49 |
| **Average FCM (ln transformed)** | | | | | |
| n = 79 individuals | **Intercept** | **8.11** | **0.16** | **50.70** | **< 0.0001** |
| | Average Docility | -0.0029 | 0.0077 | -0.38 | 0.71 |
| | Grid type | | | | |
| | high density | 0.18 | 0.10 | 1.75 | 0.082 |
| | Sex | | | | |
| | males | 0.18 | 0.11 | 1.56 | 0.12 |
